# Open source software for tube vocal tract modeling, resonance prediction, illustration, and 3D printing

**DOI:** 10.1101/2025.10.15.682256

**Authors:** Runhui Song, Jonas Beskow, Jens Edlund, Mechtild Tronnier, Rui Tu, Kecheng Zhang, Axel Ekström

## Abstract

We present accessible code-free tube vocal tract modeling software. The software implements a transfer function. Applications involve exploratory speech acoustics-based basic research and education in phonetic sciences. The program has been made publicly available and presents researchers and students in speech-centric sciences with easily accessible vocal tract modelling and vowel synthesis.

## Introduction

Tube models of the vocal tract are a historically influential method for research in speech production and acoustics (1–6). The modern iteration of this concept, with foundations in simulation through electrical engineering traces back to central work conducted by Chiba and Kajiyama (7) in Japan, Stevens (1, 8) and Fant (2, 3) in the United States and Europe. However, while the assumption is largely intuitive, its implementation has often been by engineers alone. This work is concerned with teaching the elemental aspects of speech acoustics to an audience largely unfamiliar with computational work. Generally, computational phonetics is performed by engineers (2, 3, 9) and as such place significant restrictions on participation by students in general linguistics. Here, we introduce *TubeN* – an interactable graphical user interface allowing easy simulation of tube vocal tract models and their acoustic properties. The ultimate goal of this endavour is to invite a greater number of students from general linguistics to participate in the design and execution of computational speech–centric work, and to make available to broader audiences the intricasies of speech acoustics.

### Simulating the behavior of an acoustic tube

*TubeN* implements an algorithm developed by Liljencrants and Fant (3).^1^ The algorithm calculates the formant frequencies for an acoustic tube with *M* cylindrical segments, characterized by input parameters length *L*_*n*_ and cross-sectional area *A*_*n*_. It then recursively computes a determinant through tube segments.^2^ the value of which is called *the transfer determinant* Δ_*n*_, reflecting the impedance transformation up to the *n*_*th*_ tube segment. The angular frequency *ω*, measured in radians per second (*rad/s*) is defined as:

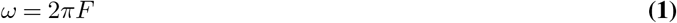

where *F* represents the frequency of the sound wave measured in Hertz (*Hz*), which corresponds to the specific acoustic frequency being simulated. For example, if the response of the tube to a 500 *Hz* sound wave is studied, then *F* = 500 *Hz*. The normalized phase angle of the *n*_*th*_ tube segment is:

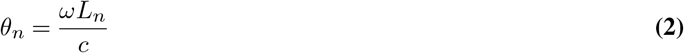

where *c* = 35300 cm/s is the speed of sound at 35*°C*, and *L*_*n*_ is the length of the *n*_*th*_ segment. The ratio of the area of two connected tube segments (*A*_*n*+1_ *and A*_*n*_) is represented as:

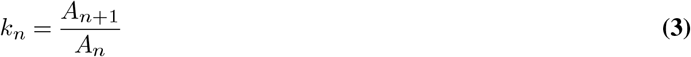

The recursive formula for the transfer determinant is:

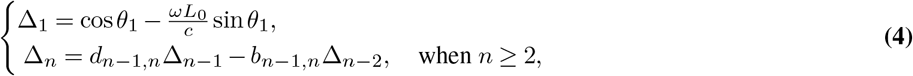

where:

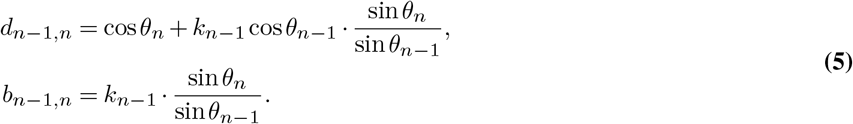

After obtaining the determinant of the final tube segment Δ_*M*_, a quasi-spectral function is constructed:

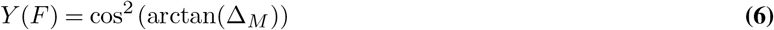

**Fig. 1.**
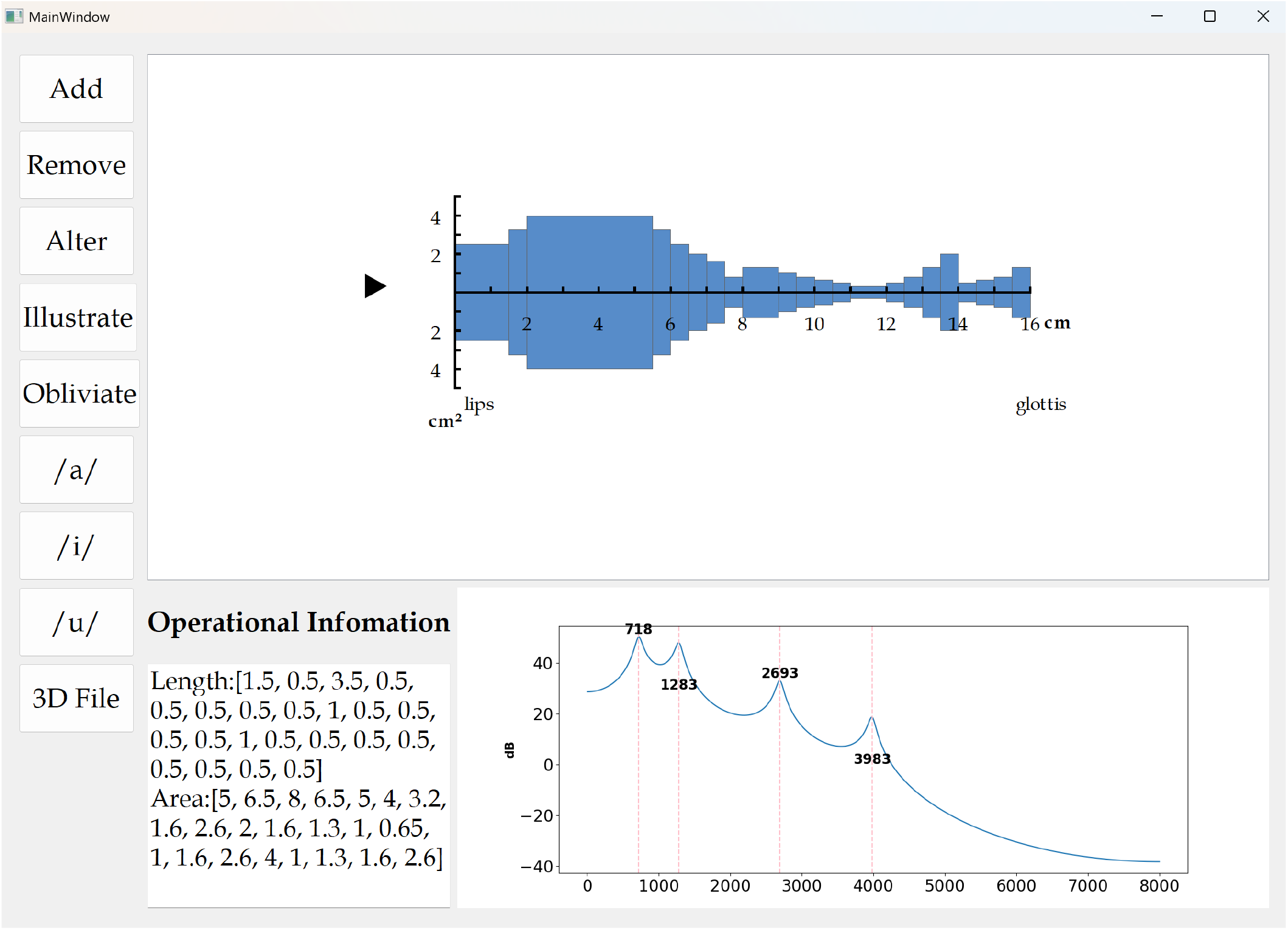
Note that where two subsequent segments are of the same area, they are rendered as a single joined segment.

### Functionalities of the graphical user interface

The software will predict formants for any sequence, and the GUI will automatically update the displayed predicted formants F1-F4 upon any change to the sequence.

#### Adding, removing and altering segments

**The *Add* function** adds *n* segments by entering their length *l* and area *a* in the following format:

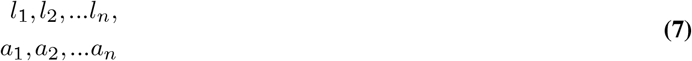

The order of input is lips to glottis, such that *l*_*1*_ corresponds to the anteriormost section (i.e., lip opening).

**The *Remove* function** lets the user remove any section in the tube sequence. It is operated by first double-clicking the relevant segment, and pressing the *Remove* button on the GUI.

**The *Alter* function** lets a user edit the length and area of any one segment after initial input. It is operated similarly to the *Remove* function, by first double-clicking the relevant segment, and pressing the *Alter* button on the GUI.

Importantly, however, segments can also be edited manually, by clicking a segment, and changing its length (by pressing the *Left arrow key* ← (to shorten) and *Right arrow key* →, (to elongate) respectively. The area *a* of the *n*th segment can be similarly altered using the Up arrow key ↑ (to expand) and Down arrow key ↓ (to contract). These “steps” are executed in increments of 0.1 cm (for length *l*) and 0.1 cm^2^ (for area *a*). For each such increment, the formant prediction window is updated accordingly, allowing for up-close inspection to changes. Finally, pressing the [TAB] key selects the segment to the right. If not segment it selected, pressing [TAB] will select the left-most segment.

**The *Obliviate* function** resets the model, and intializes a blank sheet.

**Fig. 2.**
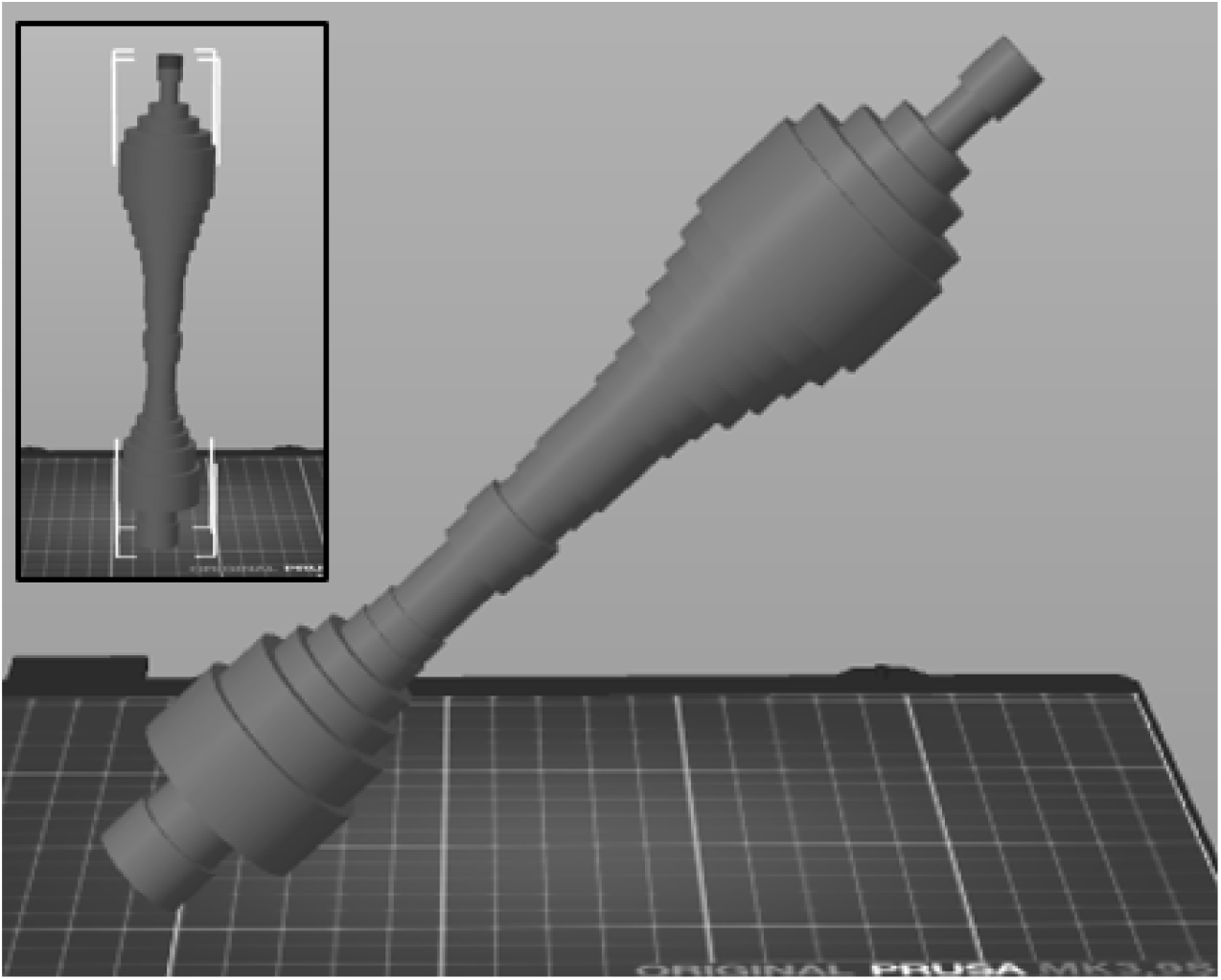
TubeN-generated 3D vocal tract. Shape is identical to that in Figure 1.

**Reference tube models** corresponding to the corner vowels [a i u], are available with single-button presses. These shapes are as reported by Fant (1971) for an adult male Russian speaker.

#### Synthesis

**A simplistic vowel synthesis function** is implemented, allowing quick-and-easy user evaluation of predicted vowel qualities. A small triangular button is placed to the left of the tube model, and users can press it to play the synthesized sound. At present, the synthesized vowel is kept at a length of 1 second, with a constant fundamental frequency of 100 Hz. More variable vowel synthesis methods are slated for implementation in future iterations.

#### Illustration

**The *Illustrate* function** allows for quick generation of illustrations for tube models, transfer and peak functions. These illustrations are intended to clearly illustrate predicted formant characteristics, and can be saved to the user’s hard drive, and may potentially be used in relevant publications or teaching materials.

#### 3D Modeling and 3D Printing

**The *3D File* function** allows for simplistic 3D modeling and printing of any arbitrary tube sequence. This button creates a 3D-printable.stl file, modeled after the currently implemented tube sequence. Because different 3D printers may require additional customization, we note that users may also slice the resultant file, and customize the relevant G-code prior to printing the tube models. We primarily recommend methods made available through the Trimesh Python library^3^ for this purpose. In our example 3D printing, we used a Prusa i3 MK3.9S 3D printer, and printed tube sequences as Polyactic Acid (PLA). We found that when a stand-in voice source (e.g., a duck call) was introduced, the intended vowel quality was reliably reproduced, provided that sufficient closure was achieved at the “closed” (i.e., the segment corresponding to the “glottis”) end of the sequence. If closure was incomplete, vowel quality was predictably degraded due to leakage. Our experience is that printing a tube model normally approx. 2.5 hours.

Note that these observations are based on only the settings described. 3D printers may utilize a range of different materials, or require unique considerations. For this reason, we cannot categorically claim that our settings are universally applicable. For example, different materials may be more or less appropriate for recreating intended vowel qualities with physical models. Finally, we do not expound on additional challenges associated with the 3D printing process; however, for the sake of increasing usability, we have made our reflections available to the reader. These can be accessed in the same GitHub folder as the TubeN program itself.^4^

**Table 1.** Starting lengths and area sections in Exercise 2. Segments are arranged in order from front to back (i.e., lips to glottis).

| Segment | Length | Area |
| --- | --- | --- |
| *Two tubes* |  |  |
| 1 | 8 cm | 4 cm<sup>2</sup> |
| 2 | 8 cm | 4 cm<sup>2</sup> |
| *Four tubes* |  |  |
| 1 | 2 cm | 4 cm<sup>2</sup> |
| 2 | 6 cm | 4 cm<sup>2</sup> |
| 3 | 2 cm | 4 cm<sup>2</sup> |
| 4 | 6 cm | 4 cm<sup>2</sup> |

### Use Case: Teaching of source/filter theory

As tube vocal tract modeling is known to facilitate teaching the principles of speech acoustics to broader audiences (6), a key reason for developing the *TubeN* software was to make tube modeling more widely available to students, teachers, and researchers outside of engineering. Below, we briefly describe our experiences conducting two workshops, offered to phonetics students, with this aim.

In–person workshops took place at Lund University, Lund, Sweden, as part of two courses – one first-cycle course, and one second–cycle course (both worth 7.5 credits each in the European Credit Transfer and Accumulation System). Students in the basic-level course took part in the workshop as part of their introductions to general phonetic science. Students in the advanced-level course were course were expected to have backgrounds in linguistics and general linguistic and phonetic theory, but little to no empirical experience of speech production analysis or modeling. Student reception was generally mixed to positive. However, in particular at the undergraduate levels, students gave positive evaluations.

#### Exercise 1

In a first exercise, students were asked to convert an MRI image of a speaker’s vocal tract into two-dimensional models.^5^ Participants were informed about the age and sex of the speaker (an adult male), but not of the vowel spoken (long close back rounded vowel [u:]). Students were instructed to trace the effective vocal tract, estimate its length given provided knowledge. They were then to segment the total estimated length into equal-length segments, and roughly estimate the area function of each. Note that this procedure does not reflect mathematical considerations typically applied by researchers in executing such conversions.^6^ Rather, it was implemented as a minimalist interpretation of such a procedure, requiring minimal preparation, training, or mathematical background, while still producing appropriate matches to the intended vowel quality. The assignment was designed to simplistically emulate a common procedure in articulatory-acoustic phonetics, where in vocal tract models are explicitly modeled on real-life data (2, 11).

#### Exercise 2

In a second exercise, students explored the impact on predicted formants resulting from changing the dimensions of simplistic and arbitrary 2-tube and 4-tube sequences (1). Participants were asked to explore the effects of stricture on simplistic vocal tract models. From uniform starting positions, they were to use the “screw” function,^7^ to explore the impact of changes on formants and vowel quality. This was to demonstrate that even rough imitations of realistic vocal tract configurations are sufficient to recreate relevant vowel qualities. As such, the goal of the exercise was to introduce participants to simplified versions of “distinctive regions” theories of speech acoustics – i.e., theories of speech production that assumes disproportionate divisions of articulatory–acoustic space. Such theories include the “Distinctive regions” theory proposed by Mrayati and colleagues (12), which presupposes eight such regions. In theory, the proposed workshop paradigm may be extended to such a narrow design; however, at present stage, given the relative lack of experience of the participants, as well as the historical historical success of two-, three- and four-tube models, we opted for this more simplistic set of tasks.

#### Reflections and future iterations

A future iteration of our “workshop” setup may seek to let participants design and 3D-print their own vocal tract models, essentially reverse engineering a vowel production phenomenon from the ground up. As the printing process is lengthy, limitations on time precluded the inclusion of such an element in our original workshop design. Nonetheless, with many universities providing so-called maker spaces, such an exercise could be designed as a take-home assignment. We have argued these are promising avenues for future work on phonetics teaching.

## Concluding thoughts

Phonetics serves to bridge the sciences of acoustics and linguistics. However, students’ expectations about speech acoustics often conform to preconceptions acquired in linguistics coursework, and rarely from direct experience with the nuances of speech production, leading to particular acoustic consequences. One main reason for the lack of relevant knowledge in acoustics is related to the lack of accessibility of appropriate pedagogical tools. Here, we presented the TubeN GUI – software that is both publicly available, and which shows promise as a teaching tool of applied phonetics. With its introduction, we hope to encourage the use of speech acoustics tools in related sciences and educational programs.

## Acknowledgements

The results of this work and the tools used will be made more widely accessible through the national infrastructure Språkbanken Tal under funding from the Swedish Research Council (2023-00161_VR). AE received additional support by the Swedish Research Council (2025–00209_VR)

## Footnotes

1 For full modeling considerations, we refer to the original publications (2, 3).

2 The original algorithm works by considering the tube behavior for one frequency at a time. However, in the current implementation, calculations are parallelized.

3 https://trimesh.org/

4 https://github.com/jbeskow/tuben

5 The data was collected during a separate data collection session – a magnetic resonance imaging (MRI) single-subject case study at the Stockholm University Brain Imaging Centre, housed at Stockholm University, Stockholm, Sweden. The scanner was a Siemens Prisma 3 Tesla whole-body MRI.

6 The reader will find such conversions described elsewhere (10).

7 Note again that this function not represented by a clickable button, but is an implicit function of the GUI.

